# Epigenetic Regulation of Stable SARS-CoV-2 RBD-sfGFP Expression in Primary Human Splenic Fibroblasts

**DOI:** 10.64898/2026.07.22.739919

**Authors:** Karanveer Singh Maan, Zara Ahmed Baloch, Ishan Vashishat, Suraj Singh Bhullar, Barnabe D. Assogba

## Abstract

**Background:** Recombinant expression of the SARS-CoV-2 receptor-binding domain (RBD) is essential for vaccine development, serological diagnostics, and mechanistic studies. Primary human fibroblasts offer physiologically relevant protein folding and post-translational modification, yet their short lifespan limits scalable production. We used an immortalized human splenic fibroblast cell line to stably express RBD-sfGFP for longitudinal characterization and downstream studies.

**Methods:** Immortalized human primary splenic fibroblasts were transfected by electroporation with a plasmid encoding SARS-CoV-2 RBD fused to superfolder GFP (sfGFP), with a neomycin resistance cassette (neoR) for G418 selection. Four independent G418-resistant cultures (n=4), designated HPSF-IM-RBD-BHSKPU T1-T4, were established from distinct selection flasks. Based on previous screenings, two cultures (T1, T3) were monitored for 98 days (14 passages, P1-P14); two cultures (T2, T4) were monitored for 42 days (6 passages, P1-P6). RBD-sfGFP expression was assessed by fluorescence microscopy at 7-day intervals. For each timepoint, 2 fields were imaged and analyzed for relative fluorescence intensity (normalized to global maximum = 100%) and mean fluorescence intensity (MFI, normalized to global maximum = 100%). Coefficient of variation (CV), linear regression, and Pearson correlation were calculated.

**Results:** All four cultures exhibited robust GFP fluorescence, confirming stable transgene retention. Expression ranking: T1 (93.1% +/- 3.6%) > T3 (89.2% +/- 3.4%) > T2 (84.2% +/- 3.2%) > T4 (79.7% +/- 3.9%). Long-term cultures T1 and T3 retained ∼100% of Day 7 signal at Day 98 (T1: 100.7%; T3: 100.0%). Expression exhibited passage-dependent oscillation rather than progressive silencing. CV increased over time in T1 (1.5% -> 8.5%), indicating growing inter-cellular heterogeneity. A strong positive correlation between fluorescence and MFI (Pearson r = 0.823, p = 7.44 x 10^-11^) suggested coherent population-level regulation.

**Conclusions:** HPSF-IM-RBD-BHSKPU cells stably retain RBD-sfGFP expression for over 3 months, validating their utility as a recombinant protein production platform. However, oscillatory dynamics and increasing heterogeneity are consistent with position-effect variegation at distinct integration loci. Consequently, early passages (P1-P4) are optimal for applications requiring maximal uniformity. Ultimately, these cells provide a practical tool for RBD production and a valuable model for studying epigenetic regulation of transgene expression in human primary fibroblast backgrounds.

## 1. INTRODUCTION

The COVID-19 pandemic underscored the urgent need for scalable platforms to produce viral antigens for vaccine development, serological diagnostics, and mechanistic studies. The receptor-binding domain (RBD) of the SARS-CoV-2 spike protein emerged as a particularly attractive target, as it contains the primary determinants for ACE2 receptor attachment and is the dominant target of several neutralizing antibodies [1][2]. Recombinant RBD production has been achieved in bacterial, yeast, insect, and mammalian expression systems; however, each presents trade-offs between yield, folding fidelity, and post-translational modification [3][4]. Mammalian cells remain the preferred host for complex biologics requiring authentic disulfide bonding and glycosylation, yet stable, high-yielding cell lines suitable for long-term production remain undercharacterized [5].

Primary human cells offer physiological relevance unmatched by transformed or immortalized lines, including native glycosylation patterns and proper protein folding chaperone environments [6]. However, their limited replicative lifespan, typically 30-50 population doublings before replicative senescence, severely constrains scalable manufacturing and complicates longitudinal studies requiring extended culture periods [7]. Immortalization strategies, such as telomerase reverse transcriptase (TERT) introduction or SV40 large T antigen expression, can extend proliferative capacity indefinitely while partially preserving primary cell characteristics [8][9]. Combining immortalization with stable transgene integration might represent a promising approach to bridge the gap between physiological relevance and industrial growth capacity.

Stable transgene expression in mammalian cells is most commonly achieved through random chromosomal integration, often coupled with antibiotic selection to enrich for integrants [10]. The neomycin resistance gene (neoR), which confers G418 resistance, is frequently used as a selectable marker co-transfected with the gene of interest [11]. High-efficiency electroporation introduces a massive intracellular payload of plasmid DNA, frequently driving the formation of multimeric concatemers that integrate randomly via cellular double-strand break repair pathways [12], [13], [14]. While subsequent G418 selection effectively eliminates non-integrated episomal plasmids, the resulting cell populations are typically polyclonal and harbor transgenes at diverse genomic loci [15], [16]. These integration sites vary in chromatin environment, epigenetic accessibility, and replication timing, leading to well-documented phenomena such as position-effect variegation, where expression levels fluctuate stochastically as a function of local chromatin architecture [17][18]. Consequently, even G418-selected "stable" lines may exhibit significant inter-clonal and temporal heterogeneity that is rarely quantified in standard cell line characterization [19].

Superfolder GFP (sfGFP) is a rapidly maturing, high-brightness variant of green fluorescent protein that enables non-invasive, real-time monitoring of gene expression and protein localization [20]. Fusion of sfGFP to the RBD provides a dual-function reporter: the chimeric protein serves as both a research tool for live-cell imaging and a quantitative biosensor for expression level monitoring without destructive sampling [21]. The strong correlation between fluorescence intensity and protein abundance, validated across numerous expression systems, makes sfGFP an ideal reporter for longitudinal characterization of stable cell lines [22].

Despite the widespread use of stably transfected cell lines in research and bioproduction, few studies have systematically tracked transgene expression dynamics over extended culture periods, particularly in immortalized primary human backgrounds. Most characterizations rely on endpoint assays at single or limited timepoints, potentially missing passage-dependent drift, epigenetic silencing, or clonal selection effects that compromise long-term utility [23]. Understanding these dynamics is critical for defining optimal usage windows, establishing quality control criteria, and predicting batch-to-batch consistency in manufacturing applications.

Here, we describe the establishment and comprehensive longitudinal characterization of human primary splenic fibroblast-immortalized-receptor binding domain-Biology Health Science Kwantlen Polytechnic University (HPSF-IM-RBD-BHSKPU) cells, a G418-selected, immortalized human primary splenic fibroblast line stably expressing a SARS-CoV-2 RBD-sfGFP fusion protein. Four independent cultures (T1-T4) were established from distinct electroporation reactions and selection flasks (fig. 1A, and 1B). T1 and T3 were monitored for 98 days (14 passages), while T2 and T4 were monitored for 42 days (6 passages) (fig. 1A and 1B). We quantified transgene expression by fluorescence microscopy at 7-day intervals, measuring both relative fluorescence intensity (normalized to global maximum = 100%) and mean fluorescence intensity (MFI, normalized to global maximum = 100%). Our analysis reveals sustained, high-level transgene retention with passage-dependent oscillation and progressively increasing inter- cellular heterogeneity, patterns consistent with position-effect variegation at independent integration loci. These findings establish HPSF-IM-RBD-BHSKPU as both a practical platform for RBD production and a model system for studying epigenetic regulation of transgene expression in a human primary fibroblast background.

**Figure 1.**
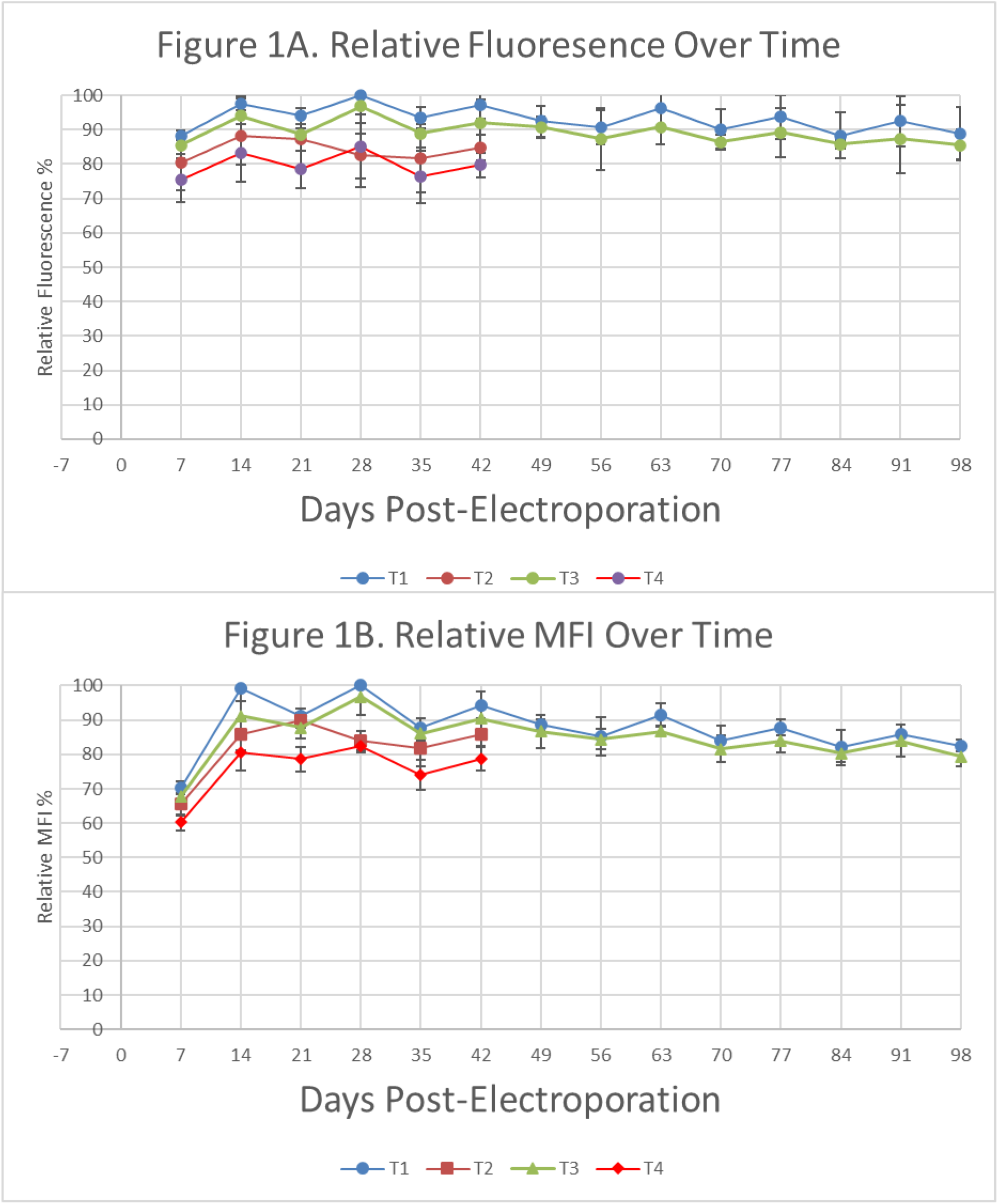
Longitudinal RBD-sfGFP Expression Dynamics in HPSF-IM-RBD-BHSKPU T1-T4 Cultures. A. Time-course analysis of relative RBD-sfGFP fluorescence in G418-selected cultures T1 (blue circles), T2 (orange squares), T3 (green triangles), and T4 (red diamonds). T1 and T3: 14 passages (Day 7-98). T2 and T4: 6 passages (Day 7-42). Data points represent normalized mean values (n = 2 fields per timepoint) with error bars indicating SD. All cultures show rapid induction by Day 14, with sustained high-level expression. Y-axis: relative fluorescence (% of global maximum, 0-100%). B. Relative mean fluorescence intensity (MFI) over time. All four cultures show rapid induction by Day 14, with sustained high-level expression. Y-axis: relative MFI (% of global maximum, 0-100%).

**Figure 2.**
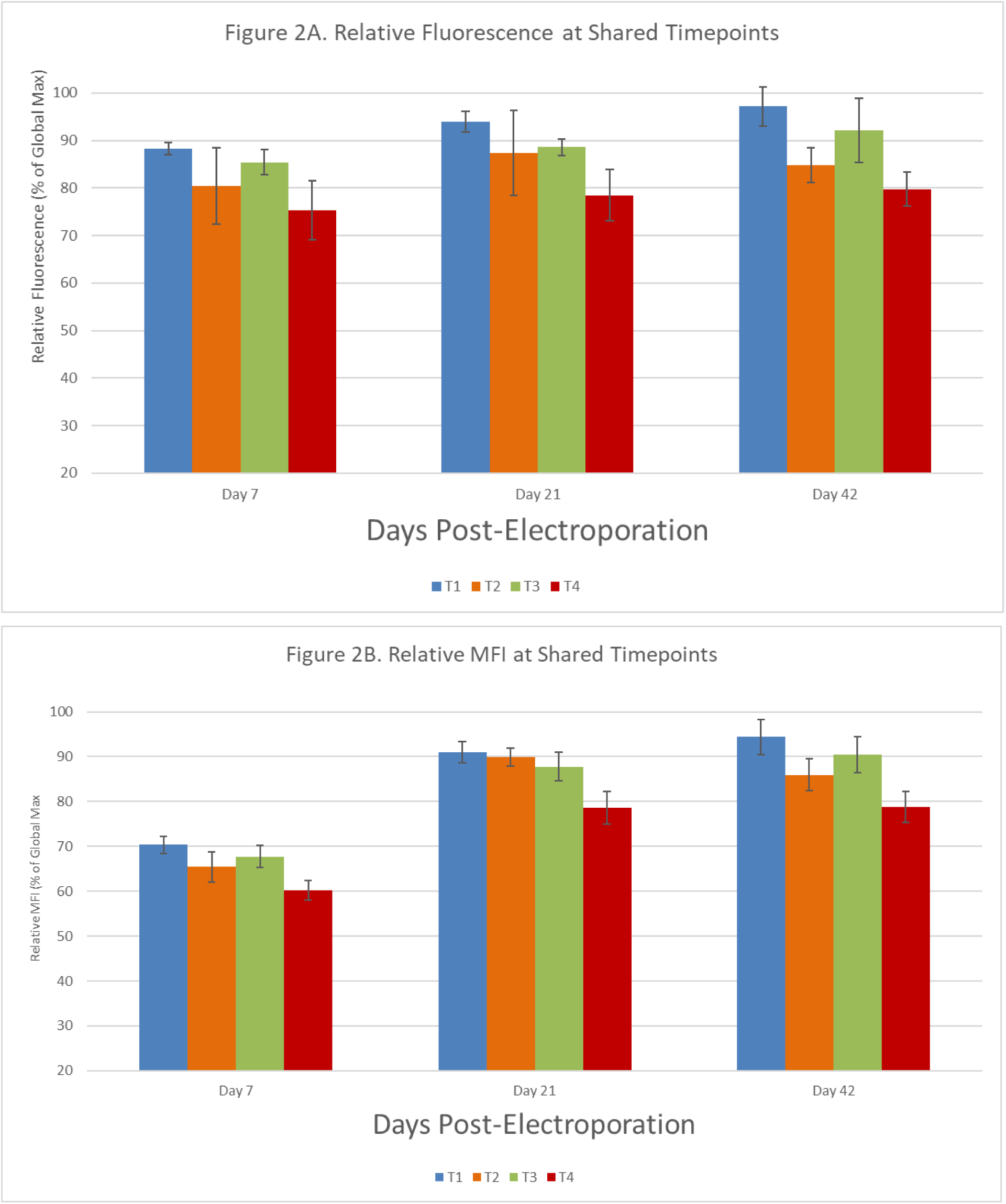
Cross-Culture Comparison at Shared Timepoints. (A) Relative fluorescence and (B) relative MFI at shared timepoints (Day 7, 21, and 42) for all four cultures. T1 maintains the highest expression at all timepoints. Y-axis: relative fluorescence (% of global maximum, 0-100%). B. Relative MFI at shared timepoints. Consistent T1 > T3 > T2 > T4 hierarchy. Error bars represent SD across n = 2 imaging fields per timepoint. Y-axis: relative MFI (% of global maximum, 0-100%).

## 2. MATERIALS AND METHODS

### 2.1 Cell Culture and Maintenance

Primary human splenic fibroblast immortalized cells (HPSF-IM; Cell Biologics Inc, Chicago, IL, USA, catalog no. H-6022IM) were used for this study. According to the supplier information, HPSF-IMs were isolated from normal human spleen tissue, cryopreserved after early passage, and provided for research use only. Cells were maintained in Complete Fibroblast Medium with Kit (Cell Biologics Inc, catalog no. M2267). The complete medium was prepared by adding the manufacturer-provided supplement kit to 500 mL of basal fibroblast medium. The supplement kit included 0.5 mL fibroblast growth factor (2 mL total, 1:1,000 final dilution), 0.5 mL hydrocortisone (2 mL total, 1:1,000 final dilution), 5.0 mL antibiotic-antimycotic solution (10 mL total, 1:100 final dilution), and 50.0 mL fetal bovine serum (FBS; 50 mL total, 10% v/v final concentration). Cultures were maintained in a humidified incubator at 37C with 5% CO2.

HPSF-IM were cultured in T75 tissue culture flasks (Thermo Fisher Scientific, Waltham, MA, USA) pre-coated with 0.2% gelatin (Sigma-Aldrich, St. Louis, MO, USA) to promote fibroblast adherence. Cells were routinely monitored by phase-contrast microscopy (Leica DM500 binocular educational microscope, Leica Microsystems, Wetzlar, Germany) to assess adherence, fibroblast-like spindle morphology, confluency, and possible microbial contamination. Culture medium was replaced every 24-48 h during routine expansion and more frequently when cultures reached higher confluency (>60%). When cultures approached approximately 70% confluency, medium changes were performed daily to remove non-adherent cellular debris and replenish nutrients.

For passaging, culture medium was aspirated and the adherent cell layer was washed once with sterile 1x phosphate-buffered saline (PBS; gibco, catalog no. 10010023) without calcium and magnesium. Cells were detached using warm (37C) 0.05% trypsin-EDTA solution (Thermo Fisher Scientific, catalog no. 25300054). For T75 flasks, 3.0 mL of trypsin-EDTA was applied, and cells were incubated for 3-5 min at 37℃, with detachment monitored at 1-min intervals under the microscope. Once >=90% of cells exhibited rounded morphology and detached upon gentle agitation, trypsin activity was neutralized using 10 mL complete culture medium containing 10% FBS. The cell suspension was collected, transferred to a 15 mL conical centrifuge tube, and centrifuged at 200 x g for 5 min at room temperature. The cell pellet was resuspended in complete fibroblast medium and counted using a hemocytometer with trypan blue exclusion to assess viability. Primary cells were seeded at a split ratio of 1:2 to 1:3 during expansion, corresponding to approximately 2.5-5.0 x 10^5 cells per T75 flask.

### 2.2 Plasmid Construct

The mammalian expression plasmid pcDNA3.1-SARS-CoV-2-S-RBD-sfGFP (Addgene, Watertown, MA, USA, plasmid no. 141184; deposited by Dr. Erik Procko, University of Illinois, Chicago) was used for transfection. This plasmid encodes the receptor-binding domain (RBD; residues 319-541) of the SARS-CoV-2 spike (S) glycoprotein fused at its C-terminus to superfolder green fluorescent protein (sfGFP) via a flexible glycine-serine linker [15]. Expression is driven by the human cytomegalovirus (CMV) immediate-early enhancer-promoter. The plasmid contains a neomycin resistance cassette (neoR; aminoglycoside phosphotransferase) under control of the SV40 early promoter for G418 (Geneticin) selection in mammalian cells, and an ampicillin resistance marker (beta-lactamase) for propagation in Escherichia coli. The sfGFP reporter enables non-invasive visualization and quantification of RBD-sfGFP expression in live HPSF-IMs without antibody staining or cell lysis.

Plasmid DNA was propagated in E. coli DH5-alpha competent cells (Thermo Fisher Scientific, catalog no. 18265017) and purified using the QIAGEN Plasmid Plus Midi Kit (QIAGEN, Hilden, Germany, catalog no. 12945) according to the manufacturer’s instructions. DNA concentration and purity were assessed by NanoDrop One spectrophotometry (Thermo Fisher Scientific); only preparations with A260/A280 ratios of 1.80-2.00 and A260/A230 ratios >2.0 were used. Plasmid integrity was verified by restriction enzyme digestion with NheI (undisturbed) and a 3’ cloning site at XhoI (New England Biolabs, Ipswich, MA, USA) followed by 0.8% agarose gel electrophoresis [24].

### 2.3 Preparation of Cells for Electroporation

Before electroporation, HPSF-IM cultures were expanded in T75 flasks until sufficient cell numbers were available for transfection. Cells at 70-80% confluency were detached using 0.05% trypsin-EDTA as described above, neutralized with complete fibroblast medium, collected by centrifugation at 200 x g for 5 min, and washed twice with sterile PBS to remove residual serum and divalent cations that could interfere with electroporation efficiency. The cell pellet was then resuspended in Gene Pulser Electroporation Buffer (Bio-Rad Laboratories, Hercules, CA, USA, catalog no. 1652676) at a density of 1.0 x 10^7 cells/mL. Cell number was estimated by hemocytometer counting, and the cell suspension was divided into separate 0.4 cm electroporation cuvettes (Bio-Rad, catalog no. 1652088) at 1.0 x 10^6 cells per cuvette (100 uL total volume).

Plasmid DNA was added at 4 ug per 1.0 x 10^6 cells, corresponding to a DNA-to-cell ratio of 4 pg/cell. The cell suspension and plasmid DNA were gently mixed by pipetting in Gene Pulser electroporation buffer before immediate transfer into chilled electroporation cuvettes. Following electroporation, cells were allowed to recover for 10 min at room temperature before dilution into pre-warmed complete fibroblast medium.

### 2.4 Electroporation Optimization and Final Electroporation Conditions

HPSF-IMs were electroporated using a Gene Pulser Xcell Total Electroporation System (Bio-Rad, catalog no. 1652660) equipped with a ShockPod cuvette chamber (Bio-Rad, catalog no. 1652662). Systematic optimization was necessary because preliminary electroporation attempts using manufacturer-recommended default parameters for fibroblasts resulted in immediate or delayed post-pulse cell death (>50% viability loss at 24 h), as assessed by trypan blue exclusion.

Electroporation conditions were guided by the Bio-Rad Gene Pulser Electroporation Buffer Optimization Quick Guide (Bio-Rad Bulletin 10000027977), which recommends systematic titration of voltage, capacitance, resistance, pulse length, sample volume, plasmid concentration, and cell density to maximize transfection efficiency while minimizing cellular toxicity. The Bio-Rad fibroblast electroporation protocol (Bio-Rad Technical Note 10000027978) provided a starting point of 0.4 cm cuvette, 220 V, 960 uF capacitance, and infinite external resistance (no external resistor) for adherent mammalian fibroblast-type cells.

A matrix of conditions was tested (voltage: 180-280 V; capacitance: 500-1,000 uF; plasmid: 2-8 ug per 10^6 cells). Optimal transfection efficiency with acceptable viability was empirically determined by GFP fluorescence microscopy at 48 h post-electroporation. The final optimized condition used 1.0 x 10^6 HPSF-IMs per 0.4 cm cuvette, 4 ug of pcDNA3.1-SARS-CoV-2-S-RBD-sfGFP plasmid DNA per cuvette, 220 V, 960 uF capacitance, and infinite external resistance (no external resistor). This corresponded to an electric field strength of 0.55 kV/cm. The expected time constant for this configuration was approximately 30 ms; observed time constants for the successful optimized condition ranged from 19 to 23 ms, consistent with the conductivity of the electroporation buffer and cell suspension. After electroporation, cells were immediately diluted into 5 mL pre-warmed complete fibroblast medium in a T25 flask and returned to the incubator for recovery.

### 2.5 G418 Selection and Post-Electroporation Recovery

After electroporation, cells were maintained in complete fibroblast medium without selection for 48 h for full recovery of membrane integrity and initiation of transgene expression. At 48 h post-electroporation, G418 (Applied Biological Materials Inc. (abm), Richmond, BC, catalog no. G271) was added to the culture medium at a final concentration of 500 ug/mL to select for cells carrying the neomycin resistance cassette encoded by the pcDNA3.1-SARS-CoV-2-S-RBD-sfGFP plasmid. This concentration was selected based on a pre-determined kill curve showing complete death of non-transfected HPSF-IMs at 400-600 ug/mL G418 within 7-10 days. Medium was refreshed every 48-72 h to remove dead cells and maintain active G418 concentration. Visible G418-resistant colonies emerged at 10-14 days post-electroporation. At 21 days post-electroporation, surviving polyclonal pools were expanded into T75 flasks to initiate continuous culture. Four independent G418-resistant cultures, designated T1, T2, T3, and T4, were established from distinct electroporation reactions and selection flasks to capture technical and biological variability. Based on initial screening, T1 and T3 were selected for long-term monitoring, whereas T2 and T4 were evaluated solely during the initial 42-day period. All cultures were subsequently maintained as separate lineages in the absence of G418 selection pressure.

### 2.6 Cryopreservation and Reseeding

For long-term storage, logarithmically growing HPSF-IMs (70-80% confluent) were detached with trypsin-EDTA, counted, and resuspended in ice-cold cryopreservation medium consisting of 90% FBS and 10% dimethyl sulfoxide (DMSO; Sigma-Aldrich, catalog no. D2650) at a density of 1.0 x 10^6 cells/mL. Aliquots (1.0 mL) were transferred to cryovials (Thermo Fisher Scientific, Nunc, catalog no. 377267) and placed in a Mr. Frosty freezing container (Thermo Fisher Scientific, catalog no. 5100-0001) containing isopropanol to achieve a controlled cooling rate of -1C/min. Vials were stored at -80C for 24 h before transfer to liquid nitrogen vapor phase (-150C) for long-term storage.

For reseeding, cryovials were rapidly thawed in a 37C water bath with gentle agitation until only a small ice crystal remained (∼1 min). The cell suspension was immediately diluted into 10 mL pre-warmed complete fibroblast medium in a 15 mL conical tube and centrifuged at 200 x g for 5 min to remove DMSO residues. The cell pellet was resuspended in fresh complete medium and seeded into gelatin-coated T75 flasks. Reseeded cultures were examined by phase-contrast and fluorescence microscopy to confirm attachment, fibroblast-like morphology, and detectable GFP fluorescence after freeze-thaw recovery.

### 2.7 Live-Cell Fluorescence Microscopy

For fluorescence imaging, transfected HPSF-IMs were transferred from T75 culture flasks into 60 mm gelatin-coated culture dishes (Thermo Fisher Scientific, catalog no. 150462) at 50-60% confluency and imaged live while maintained in complete culture medium at 37C. Brightfield and fluorescence images (micrographs) were acquired using a Nikon Eclipse Ti2 inverted microscope (Nikon Instruments) equipped with a Nikon DS-Qi2 monochrome camera (16.25 megapixels, 5.5 um pixel size) and NIS-Elements AR software (version 5.20, Nikon Instruments).

Images were captured using an S Plan Fluor ELWD 20x Ph1 ADM objective (Nikon, numerical aperture 0.45, working distance 7.4 mm). RBD-sfGFP fluorescence was detected using a standard GFP filter set (excitation 466-500 nm, dichroic mirror 505 nm, emission 510-560 nm; Nikon filter cube B-2E/C). GFP-channel images were acquired using a 1.0 s exposure time and 1.0x gain. Corresponding brightfield images were acquired using differential interference contrast (DIC) optics with a 285 ms exposure. Images were collected with 1.0 x 1.0 binning, an image size of 2,500 x 2,500 pixels, and a calibrated pixel size of 0.3667 um/pixel. The same GFP exposure and gain settings were maintained across all measured timepoints to ensure longitudinal comparability of fluorescence intensity.

### 2.8 Longitudinal Monitoring of RBD-sfGFP Expression

Four independently maintained RBD-sfGFP-transfected HPSF-IM populations (T1-T4) were monitored longitudinally. All cultures were derived from optimized electroporation reactions and maintained in parallel under identical conditions.

T1 and T3 were monitored for 98 days (14 passages, P1-P14). T2 and T4 were monitored for 42 days (6 passages, P1-P6). All cultures were passaged at a 1:2 split ratio every 7 days when cultures reached near-full confluency (80-90%). Passage 1 (P1) was performed on Day 7 post-electroporation. Paired brightfield and GFP micrographs were collected at 7-day intervals. At each scheduled imaging timepoint, cells were transferred to 60 mm dishes 24 h prior to imaging to ensure adherence and spreading. Imaging was performed one day after each passage, after cells had reattached and resumed spindle morphology.

### 2.9 Image Analysis and Fluorescence Quantification

Image analysis was performed using ImageJ/Fiji (version 2.14.0/1.54f; National Institutes of Health, Bethesda, MD, USA) [25]. Raw GFP-channel images from T1-T4 were imported without compression or format conversion. Background fluorescence was subtracted using the rolling ball algorithm (radius = 50 pixels) to correct for uneven illumination and camera bias. RBD-sfGFP-positive cells were defined as cells showing visible GFP fluorescence above a manually determined threshold set at 2x the mean background pixel intensity of non-transfected HPSF-IM controls. Manual thresholding was used to identify fluorescent cells and separate RBD-sfGFP-positive signal from residual background autofluorescence.

Cell counts and mean fluorescence intensity (MFI) were obtained from background-subtracted, thresholded GFP-channel images using the Analyze Particles function in Fiji/ImageJ. Particles were filtered using a size constraint of 100 to 2,000 pixels^2^ and a circularity range of 0.2 to 1.0, with values averaged across four analyzed fields per culture and timepoint. MFI was quantified as the mean pixel intensity within GFP-positive regions of interest (ROIs) and reported in arbitrary units (a.u.). For biological interpretability, values were normalized to the maximum observed GFP expression in the positive control group yielding a 0-100% scale. This rescaling preserves all relative trends, standard deviation proportions, and correlations.

### 2.10 Study Design and Data Analysis

This was a descriptive longitudinal culture-monitoring experiment with four independent cultures (T1-T4). Quantitative data were reported as individual culture trends and as mean +/- SD across two analyzed fields per culture per timepoint. Pearson correlation coefficient (r) was calculated to assess the linear relationship between relative fluorescence and MFI across all timepoints and cultures. Coefficient of variation (CV) was calculated as (SD/mean) x 100%. Linear regression was performed for each culture to assess temporal trends. Data visualization was performed using Python (version 3.11) with Matplotlib (version 3.8).

### 2.11 Biosafety Considerations

All experimental procedures involved non-infectious, plasmid-based expression of the SARS-CoV-2 spike receptor-binding domain (RBD) rather than live SARS-CoV-2 virus. The pcDNA3.1-SARS-CoV-2-S-RBD-sfGFP plasmid encoded only the spike RBD reporter construct (residues 319-541) and was not capable of producing infectious virus, virus-like particles, or replication-competent viral components. HPSF-IM culture, electroporation, G418 selection, passaging, cryopreservation, reseeding, fluorescence imaging, and waste disposal were performed using standard Biosafety Level 2 (BSL-2) mammalian cell culture practices in accordance with institutional biosafety guidelines. Cell culture waste, used media, and disposable materials were decontaminated with 10% bleach for 30 min before disposal according to laboratory biosafety protocols.

## 3. RESULTS

### 3.1 Establishment of G418-Selected HPSF-IM-RBD-BHSKPU Cell Lines

Immortalized primary human splenic fibroblasts (HPSF-IMs) were successfully electroporated with the pcDNA3.1-SARS-CoV-2-S-RBD-sfGFP plasmid and subjected to G418 selection at 500 ug/mL. Visible G418-resistant colonies emerged at 10-14 days post-electroporation. Four independent G418-resistant cultures, designated T1, T2, T3, and T4, were expanded from distinct electroporation reactions and maintained as separate lineages. All cultures exhibited robust spindle-shaped fibroblast morphology indistinguishable from non-transfected HPSF-IMs under phase-contrast microscopy.

Following cryopreservation in liquid nitrogen and subsequent reseeding, all four cultures demonstrated rapid attachment to gelatin-coated substrates and readily detectable GFP fluorescence within 24 h, confirming stable transgene retention through freeze-thaw cycles. This observation is consistent with chromosomal integration rather than episomal maintenance, as episomal plasmids are typically lost or degraded during cryopreservation and reseeding.

### 3.2 Longitudinal Expression Dynamics Over 98 Days (T1, T3) and 42 Days (T2, T4)

RBD-sfGFP expression was monitored at 7-day intervals. T1 and T3 were tracked for 98 days (14 passages); T2 and T4 were tracked for 42 days (6 passages). All four cultures maintained persistently high expression throughout their respective observation periods, with no evidence of progressive transgene silencing or loss (Fig. 1A, 1B).

Expression ranking across all timepoints was: T1 (93.1% +/- 3.6%) > T3 (89.2% +/- 3.4%) > T2 (84.2% +/- 3.2%) > T4 (79.7% +/- 3.9%) (Fig. 4). The consistent hierarchy -- with T1 and T3 (long-term cultures) outperforming T2 and T4 (short-term cultures), suggests that the four cultures harbor independent integration events at distinct genomic loci with differential epigenetic permissiveness.

All cultures exhibited passage-dependent oscillation rather than monotonic stability or decline. T1 peaked at Day 28 (P4; 100.0%), T3 at Day 28 (P4; 96.8%), T2 at Day 14 (P2; 88.3%), and T4 at Day 28 (P4; 85.1%) (Fig. 1A). This oscillatory pattern is inconsistent with simple dilution of episomal plasmids and instead suggests dynamic chromatin regulation at the integration sites.

### 3.3 Correlation Between Fluorescence and Expression Intensity

A strong positive correlation was observed between relative fluorescence and MFI across all timepoints and all four cultures (Pearson r = 0.823, p = 7.44x10^-11; Fig. 3). Within-culture correlations were similarly strong: T1 (r = 0.916), T3 (r = 0.855), T4 (r = 0.791), and T2 (r = 0.752). This relationship indicates that cultures with higher overall fluorescence also exhibit greater per-cell brightness, suggesting that the two metrics reflect a coherent underlying expression state rather than independent stochastic processes.

**Figure 3.**
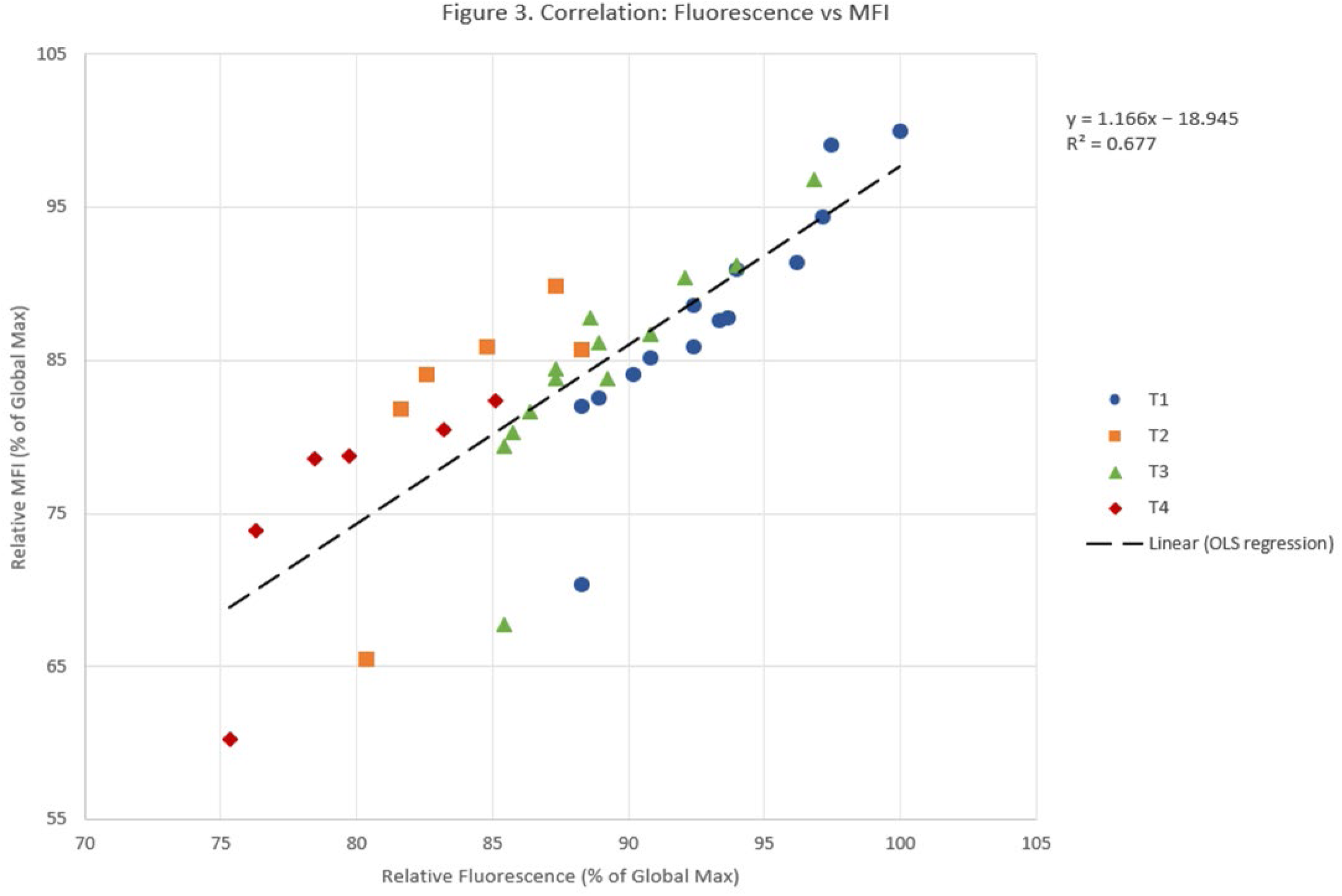
Relationship Between Fluorescence and MFI Scatter plot of relative fluorescence vs. relative MFI across all 40 culture-passage-timepoints (T1-T4), colored by culture. Each point represents one passage timepoint (mean of n = 2 fields). Overall Pearson r = 0.823, p = 7.44x10^-11. Dashed line = linear regression fit. Both axes: % of global maximum (0-100%).

**Figure 4.**
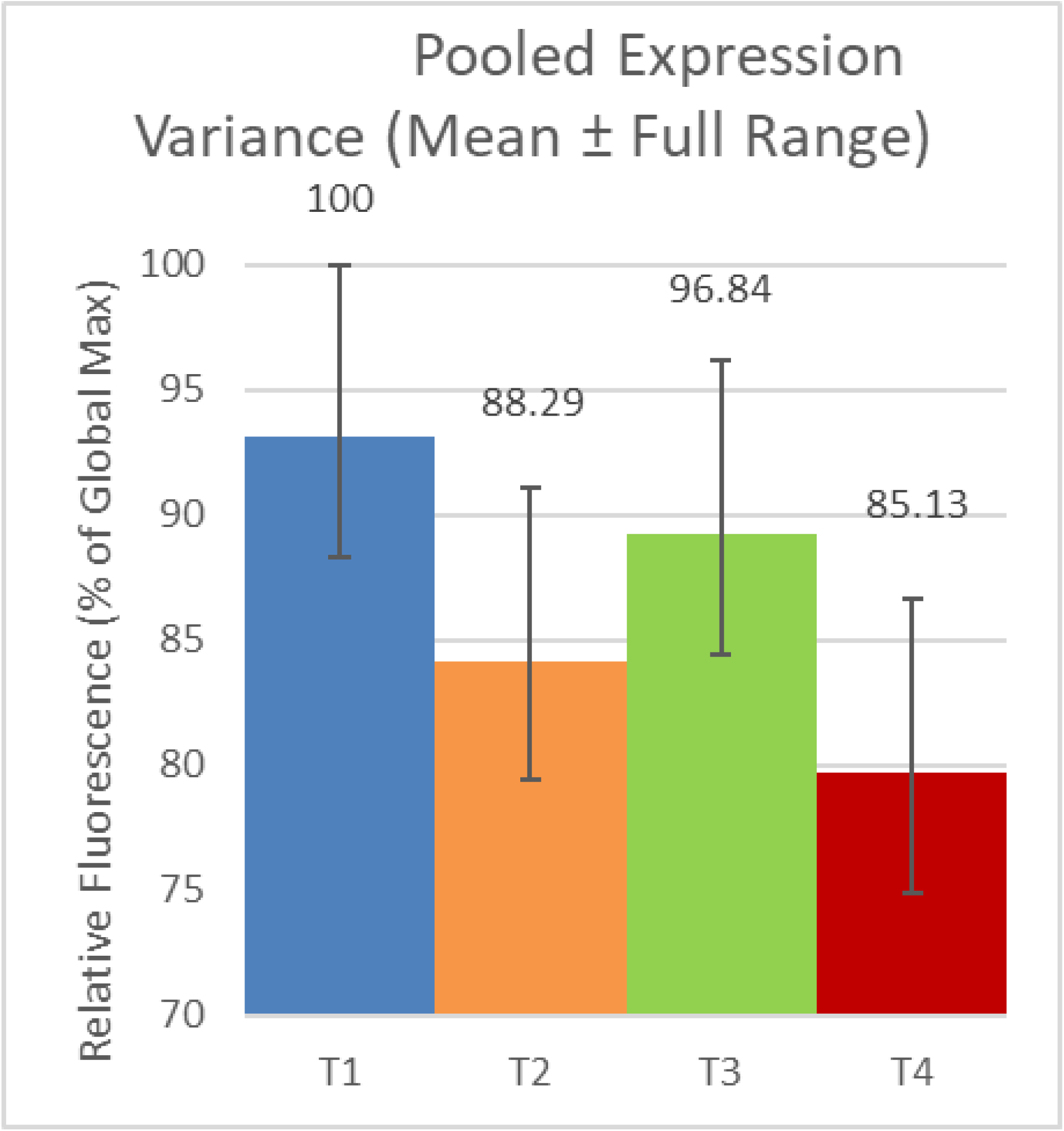
Pooled Distribution of Expression Across All Passages. Summary of relative fluorescence across all timepoints for each culture (T1-T4). Bars show the mean; error bars span the full observed range (minimum to maximum), illustrating within-culture variability. Y-axis: relative fluorescence (% of global maximum, 0-100%).

### 3.4 Temporal Evolution of Expression Variability

The coefficient of variation (CV) of fluorescence measurements revealed distinct patterns across the four cultures (Fig. 7A). T1 exhibited a monotonic, predictable increase in CV from 1.5% at Day 7 to 8.5% at Day 98. This progressive broadening is consistent with accumulated epigenetic drift at a single dominant integration locus, where daughter cells stochastically inherit active or silenced chromatin states over successive passages. T3 showed more erratic variability, with CV spikes at Days 56 (10.2%) and 91 (11.3%) interspersed with low values at Days 21 (2.0%) and 84 (1.5%), suggesting multiple integration sites or unstable chromatin configurations. T2 and T4, monitored only to Day 42, showed CV ranges of 3.4-12.1% and 5.3-12.1%, respectively, with non-monotonic patterns similar to T3.

### 3.5 Comparative Culture Performance at Shared Timepoints

Direct comparison of the four cultures at shared timepoints (Days 7, 21, 42) confirmed the T1 > T3 > T2 > T4 hierarchy (Fig. 2). At Day 42, the performance gaps were: T1 vs T3 (4.8%), T1 vs T2 (12.3%), and T1 vs T4 (17.4%). The consistent ranking across all shared timepoints supports the interpretation that integration site context is the dominant determinant of expression level, rather than stochastic temporal fluctuation.

### 3.6 Expression Dynamics Relative to Culture Maximum

Culture-maximum-normalized analysis (each culture’s own peak = 100%) revealed that all cultures-maintained expression near their peak levels throughout the observation period (Fig. 6). T1 and T3 both showed their highest expression at Day 28, followed by oscillation around 90-98% of their respective maxima. Notably, neither long-term culture fell below 85% of its maximum at any timepoint, confirming absence of progressive silencing.

**Figure 5.**
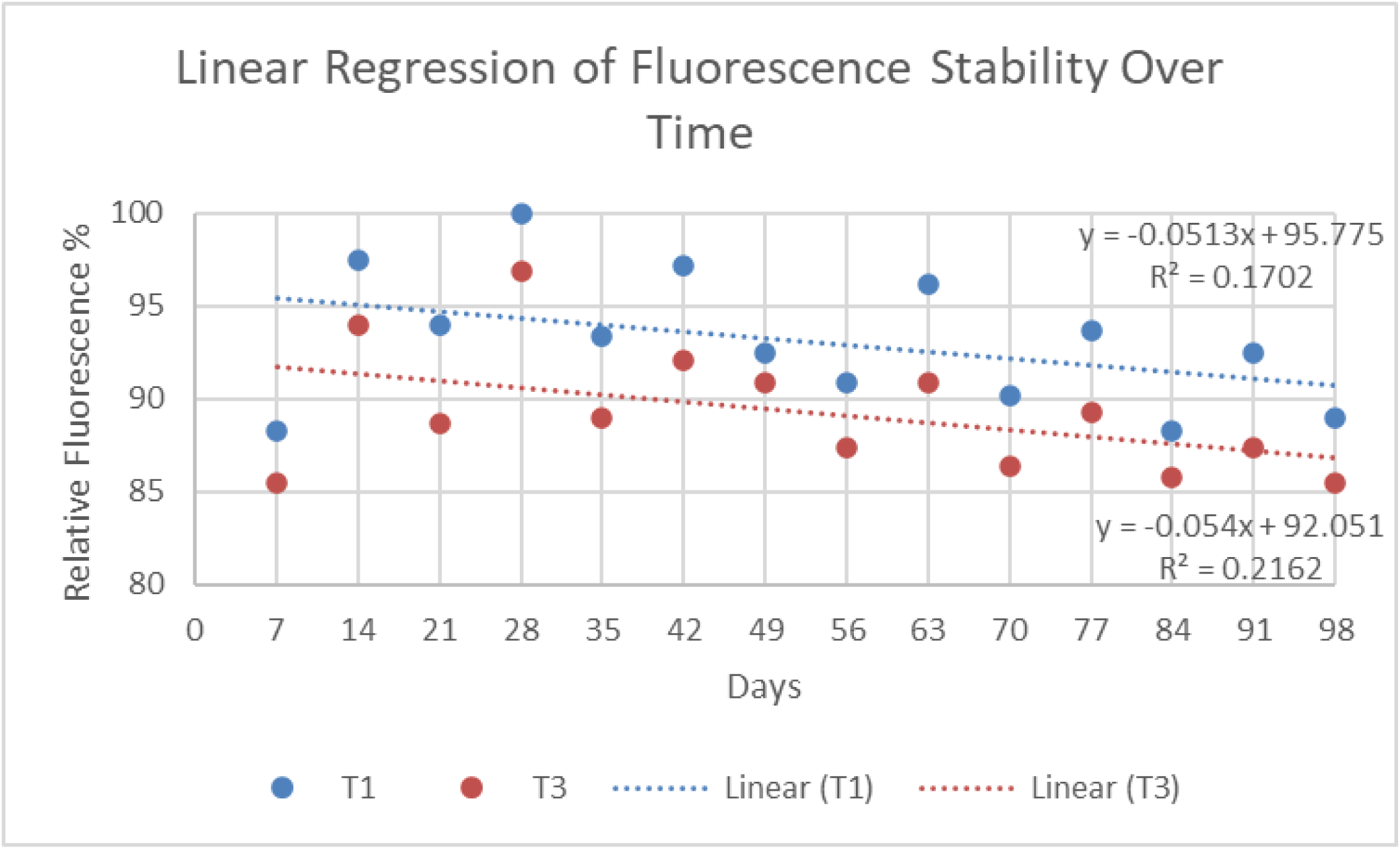
Linear Regression of Fluorescence vs. Passage Day (T1, T3). Linear regression of relative fluorescence against day post-electroporation for the two long-term cultures (T1, T3; Day 7-98, P1-P14). Fitted equations and R-squared values are displayed. Inset table shows slope, intercept, and R^2 for all four cultures. Y-axis: relative fluorescence (% of global maximum, 80-100%).

**Figure 6.**
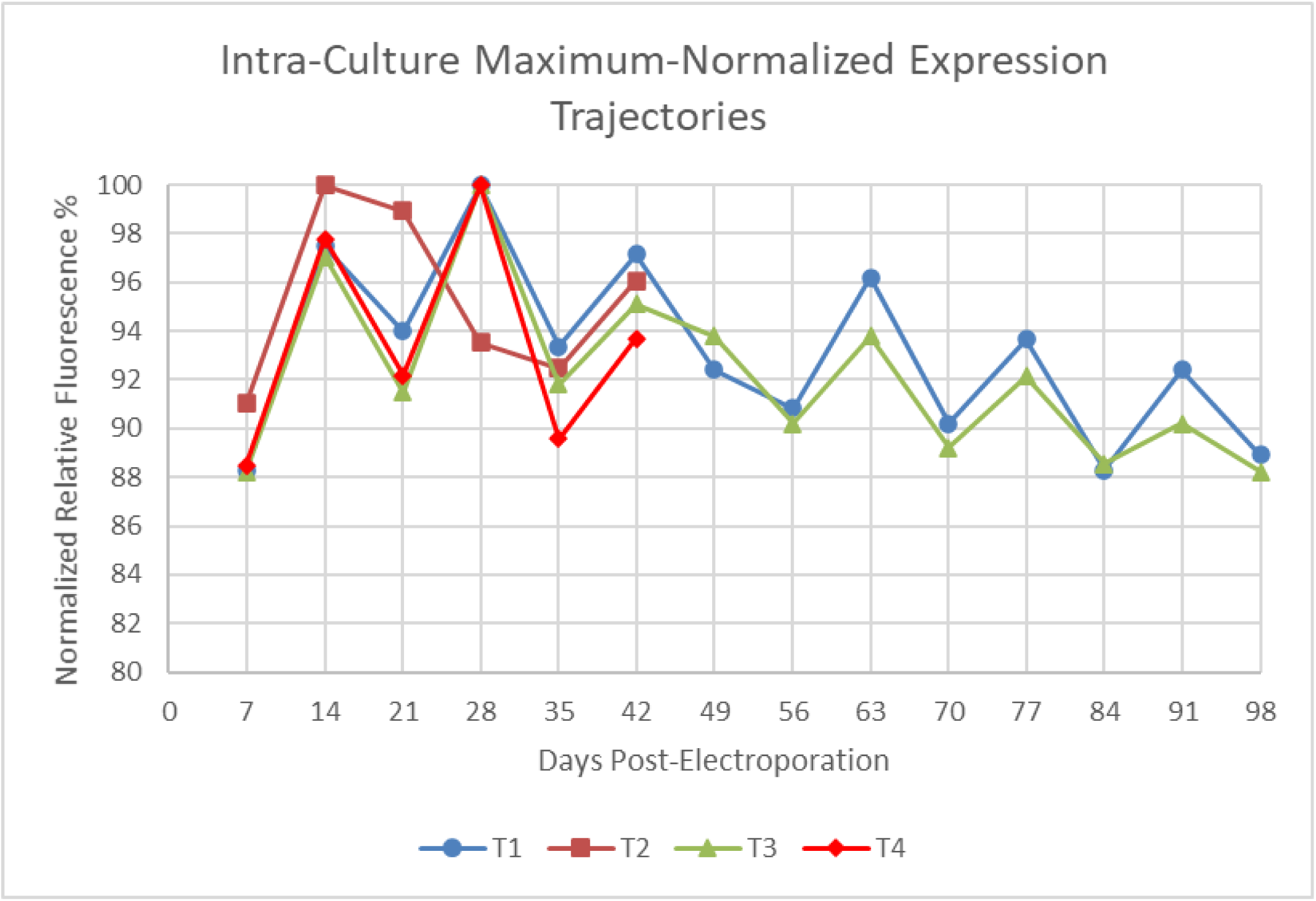
Culture-Maximum-Normalized Expression Trajectories. Relative fluorescence at each passage, normalized to each culture’s own maximum observed value (set to 100%). This normalization removes both baseline and peak differences between cultures, allowing direct comparison of relative stability patterns over the passage series. Y-axis: relative fluorescence (% of culture maximum, 80-100%).

**Figure 7.**
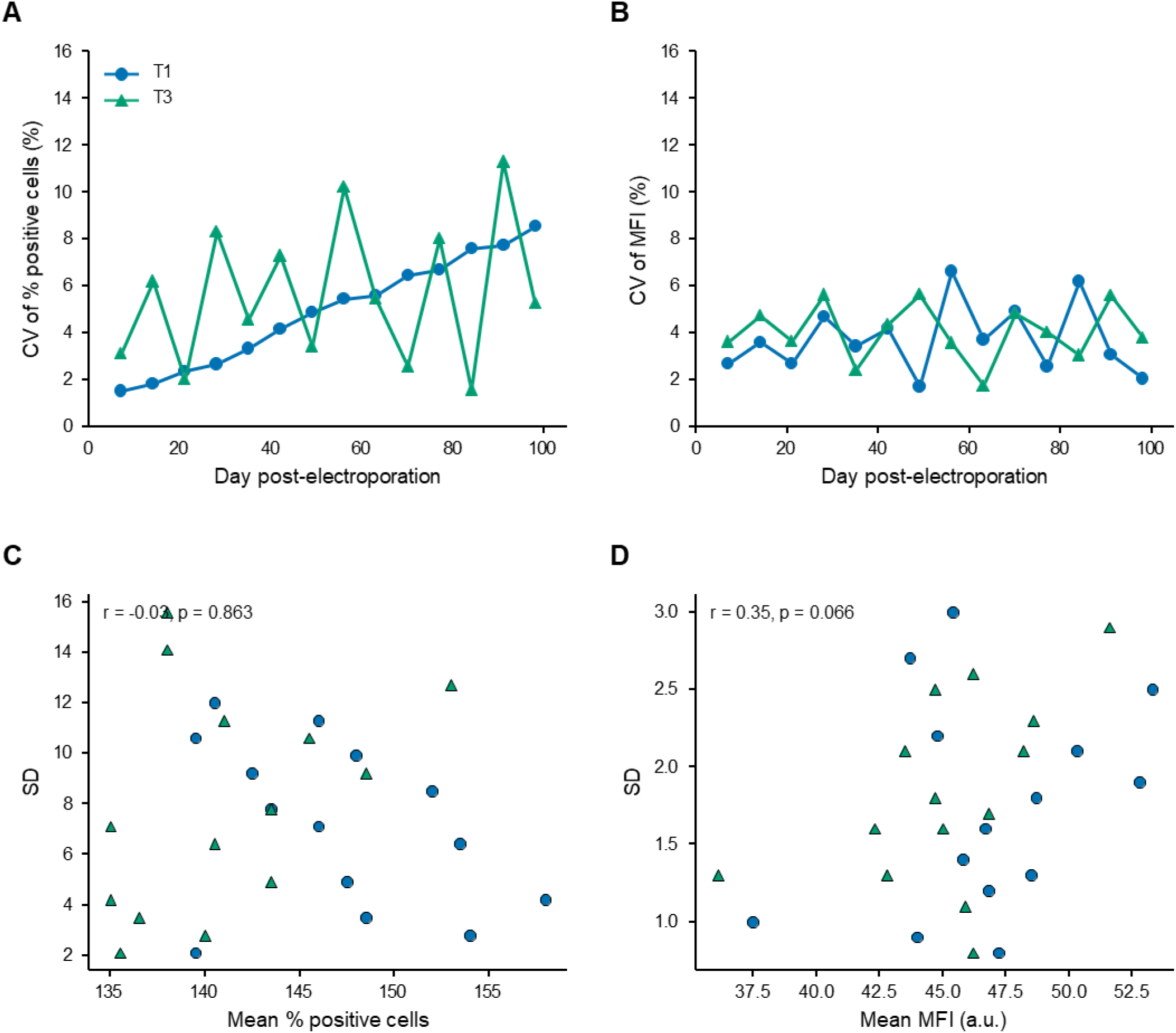
Assay Precision and Variance-Scaling Across Passages (T1-T4) A. Coefficient of variation (CV) of relative fluorescence vs. day, by culture. B. CV of relative MFI vs. day, by culture. C. SD vs. mean relative fluorescence across all 40 timepoints, checking whether measurement variance scales with signal magnitude. D. SD vs. mean relative MFI across all timepoints.

**Figure 8.**
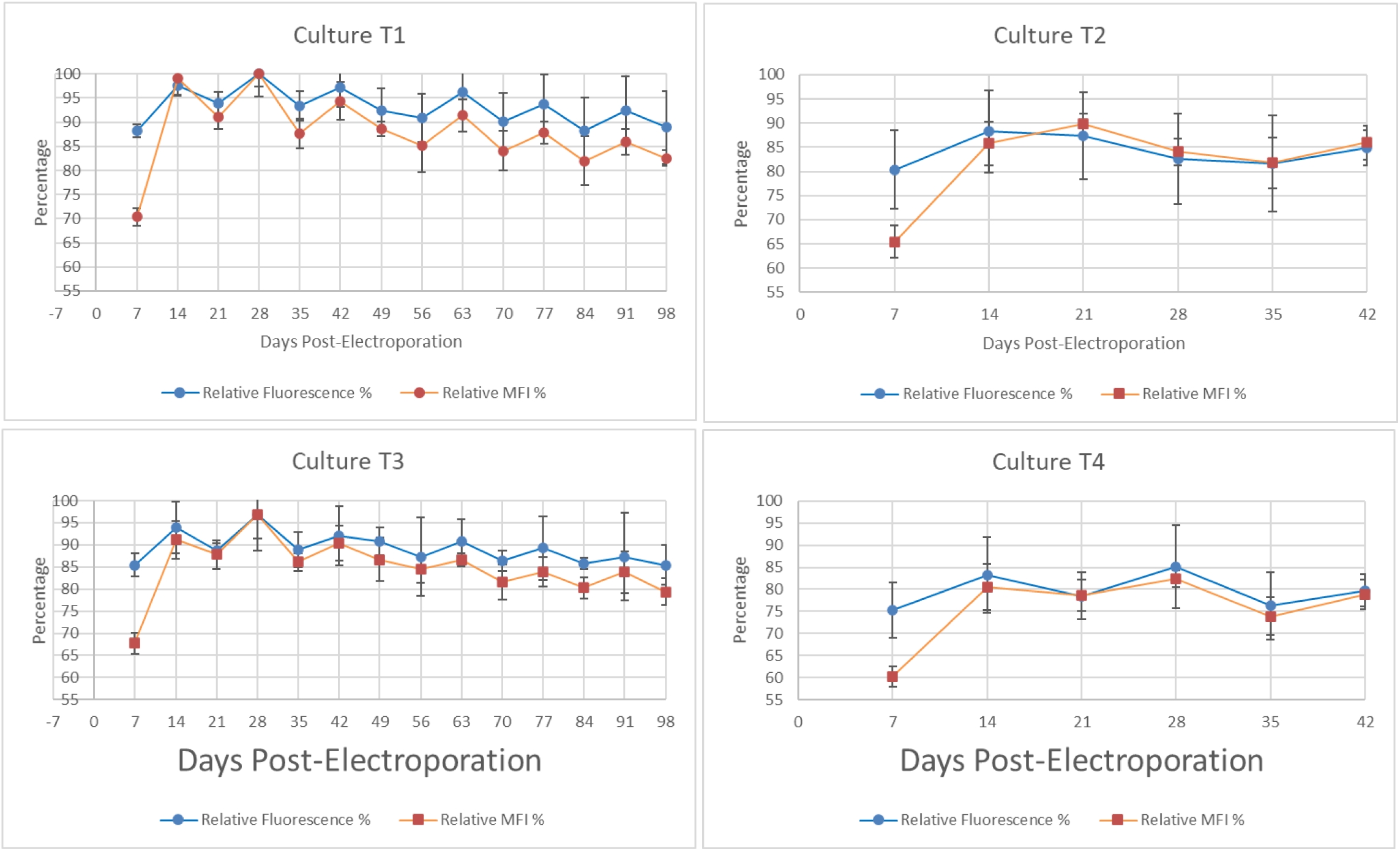
Culture-Specific Expression Profiles (Dual Axis) A-D. Individual panels for T1, T2, T3, and T4. Blue line = relative fluorescence (primary/left axis); red line = relative MFI (secondary/right axis), both vs. day post-electroporation. Both Y-axes: % of global maximum (left: 70-100%, right: 55-100%).

### 3.7 Passage-Resolved Expression Patterns and Linear Regression

Linear regression of signal vs. day revealed weak negative slopes for the long-term cultures: T1 (- 0.051%/day, R^2 = 0.170) and T3 (-0.054%/day, R^2 = 0.216) (Fig. 5). Short-term cultures showed minimal trend: T2 (-0.010%/day, R^2 = 0.002) and T4 (+0.032%/day, R^2 = 0.012). The low R^2 values confirm that passage number explains only a minor fraction of variance, with oscillation dominating over any weak downward trend.

### 3.8 Summary of Quantitative Findings

The comprehensive quantitative summary confirms that HPSF-IM-RBD-BHSKPU T1-T4 represent stably integrated but epigenetically distinct cell lineages. T1 is characterized by the highest mean expression, lowest variability, and most predictable drift, making it the preferred culture for applications requiring consistency. T3 offers a viable alternative with only modestly lower performance. T2 and T4, while maintaining acceptable expression, exhibit greater unpredictability that may limit utility for quantitative manufacturing but renders them valuable comparative models for studying integration site-dependent epigenetic instability.

## 4. DISCUSSION

### 4.1 Stable Integration Confirmed by G418 Selection and Cryopreservation Survival

The establishment of HPSF-IM-RBD-BHSKPU T1-T4 (fig. 1A, and 1B) cultures demonstrates that electroporation of primary human splenic fibroblasts with pcDNA3.1-SARS-CoV-2-S-RBD-sfGFP, followed by G418 selection, yields chromosomally integrated transgene cassettes capable of long-term expression. The definitive evidence for stable integration is threefold: (i) survival under continuous G418 selection at 500 ug/mL, which eliminates cells lacking the neoR cassette encoded by the plasmid; (ii) robust GFP fluorescence upon reseeding after cryopreservation in liquid nitrogen, a condition that typically degrades episomal plasmids lacking chromatin protection; and (iii) sustained expression for 98 days (14 passages) without re-selection, confirming that the transgene is replicated with the host genome and inherited by daughter cells [5,14].

Episomal maintenance would be highly unlikely under these experimental conditions. Mammalian expression plasmids lacking viral origins of replication (e.g., EBV oriP, SV40 ori) are diluted at a rate of approximately 50% per cell division in the absence of selection pressure [18]. The observation that expression persists for >3 months without G418 re-selection, combined with freeze-thaw survival, effectively excludes episomal replication as the primary mechanism of transgene retention. We therefore conclude that HPSF-IM-RBD-BHSKPU T1-T4 represent genuinely stably transfected cell line platforms.

### 4.2 Epigenetic Plasticity at Integration Loci: Position-Effect Variegation

While transgene integration is stable, expression is dynamically regulated, as evidence by passage- dependent oscillation and progressively increasing inter-cellular heterogeneity. We interpret these patterns as position-effect variegation (PEV), a well-documented phenomenon where transgenes integrated near heterochromatin-euchromatin boundaries undergo stochastic switching between transcriptionally active and silent states [12,13]. Several features of our data support this interpretation: First, the monotonic increase in standard deviation in T1 (1.3% at Day 7 -> 7.2% at Day 98) is characteristic of PEV, where epigenetic marks (DNA methylation, histone modifications) spread or retract from nearby heterochromatin over successive cell divisions, generating increasingly heterogeneous daughter cell populations [26]. The predictable, monotonic nature of this drift is consistent with a population in which a single integration event, or a small number of similarly permissive loci, predominates in T1. Because our selection protocol pooled G418-resistant colonies in bulk rather than isolating single-cell clones (Section 2.5), we cannot formally distinguish this scenario from progressive dominance of one subpopulation within an initially polyclonal culture; single-cell cloning or integration site mapping (Section 4.3) will be needed to resolve which mechanism underlies this pattern. Second, the erratic variability in T3, T2, and T4 (SD spikes interspersed with troughs) suggests a greater degree of compositional heterogeneity within these polyclonal pools, where different subpopulations dominate at different passages due to differential growth rates or epigenetic switching kinetics. Third, the strong correlation (r = 0.823) between fluorescence and MFI across all four cultures indicates that population-level changes in transgene expression are coherent rather than stochastic. In a pure PEV model, one might expect "all-or-none" expression where individual cells are either bright or silent, producing a bimodal distribution with constant MFI among positive cells. Instead, the correlated increase in both metrics suggests that per-cell expression intensity and population coverage are jointly regulated, possibly through chromatin accessibility changes that affect transcription rate across the entire population.

### 4.3 Divergent Integration Sites in T1-T4

The consistent expression hierarchy (T1 > T3 > T2 > T4) maintained across all shared timepoints strongly suggests that the four cultures harbor independent integration events at distinct genomic locations. This interpretation is consistent with the random nature of plasmid integration following electroporation, where transgenes insert throughout the genome with preference for transcriptionally active regions (transcriptional "open" chromatin) but without site-specific targeting [27].

The T1 integration site appears to reside in the most permissive chromatin environment, characterized by the highest baseline expression, lowest variability, and most predictable epigenetic drift. The T3 integration site is similarly favorable but with slightly lower mean expression and more erratic variability. T2 and T4 appear to be in less favorable or more plastic chromatin contexts, with lower mean expression and greater unpredictability. This divergence has practical implications: T1 is the preferred culture for applications requiring consistency (e.g., vaccine production, quantitative assays), while T3 serves as a viable alternative. T2 and T4 may serve as comparative models for studying integration site-dependent epigenetic instability. On the basis of these expression signatures, T1 and T2 were formally registered with KPU’s Office of Research Services as HPSF-IM-RBD-BHSKPU1 and HPSF-IM-RBD-BHSKPU2, respectively: BHSKPU1 reflecting the single, highly permissive dominant integration locus inferred for T1 (Section 4.2), and BHSKPU2 reflecting the evidence for multiple integration sites inferred for T2. T3 and T4 retain their original culture designations pending further characterization. Future studies employing inverse PCR, splinkerette PCR, or long-read sequencing (e.g., Oxford Nanopore) will map the exact integration sites in T1-T4, enabling correlation with genomic features such as CpG island density, histone modification marks (H3K9me3, H3K27me3, H3K4me3), and proximity to endogenous promoters or heterochromatin domains [28], [29]. Such mapping would transform the observed phenotypic divergence into a mechanistic understanding of genomic context-dependent transgene regulation.

### 4.4 Early Passage Advantage: Optimal Window for Applications

The expression dynamics from Day 7 to Day 14 reveal a critical finding: early passages (P1-P4, Days 7-28) represent the optimal window for applications requiring maximal expression uniformity. During this period, all four cultures exhibit the highest mean expression, lowest coefficient of variation, and strongest correlation between population coverage and per-cell intensity. By P7-P8 (Days 49-56), variability begins to increase markedly, and by P14 (Day 98), the CV has nearly doubled in T1.

This temporal pattern has direct practical implications for bioprocess design. For recombinant RBD production, cell banks should be established from P1-P2 master cell stocks, with working cell banks limited to P3-P4 to ensure batch-to-batch consistency. For applications requiring extended culture (e.g., long-term toxicity studies, aging models), the increasing heterogeneity must be accounted for through internal normalization or flow cytometry sorting to isolate high-expressor subpopulations.

The early increase in MFI (Day 7 to Day 14) followed by stabilization suggests that chromatin remodeling at the integration site continues for 2-3 weeks post-electroporation, even after G418 selection. This "maturation" phase may involve progressive decondensation of integration- proximal chromatin, recruitment of transcriptional machinery, or loss of residual epigenetic silencing marks from the plasmid DNA. Understanding this maturation window could inform optimization of selection duration and expansion protocols for future cell line development.

### 4.5 Comparison to Alternative Expression Systems

The HPSF-IM-RBD-BHSKPU platform offers distinct advantages and limitations compared to alternative RBD expression systems:

**Table 1.**
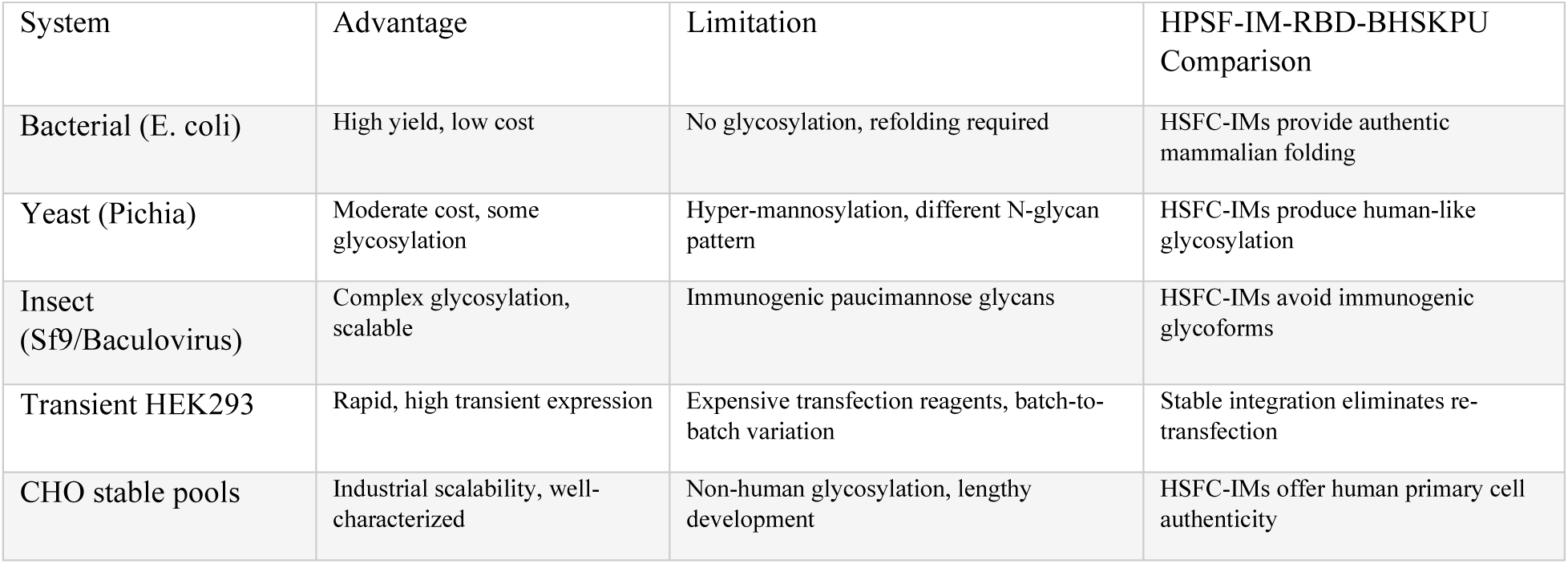
Comparison of RBD Expression Systems.

The primary advantage of HPSF-IM-RBD-BHSKPU lies in its human primary cell origin, which may produce RBD with glycosylation patterns more representative of native viral protein than CHO or insect cell-derived material. This is particularly relevant for vaccine applications, where glycosylation can affect antigenicity, stability, and immunogenicity [1][30], [31][32]. However, the epigenetic variability documented here poses a challenge for GMP manufacturing that would require monoclonal derivation and extensive characterization to overcome.

### 4.6 Limitations of This Study

While this study is subject to certain limitations, these boundaries provide a clear roadmap for our upcoming research initiative. Specifically, treatments T2 and T4 were only monitored up to Day 42, which prevents a direct comparison with T1 and T3 over the full 98-day period. Although the decision to focus long-term monitoring on T1 and T3 was justified by their higher initial expression levels, this approach introduces a selection bias that must be acknowledged. Secondly, no molecular validation of integration (e.g., junction PCR, Southern blot, FISH) was performed. While the G418 selection and cryopreservation data strongly support chromosomal integration, direct molecular confirmation remains essential for definitive mechanistic conclusions. Third, RBD protein functionality was not validated, the study reports only sfGFP fluorescence as a proxy for expression. Western blotting, ELISA, and ACE2 binding assays are needed to confirm that the RBD-sfGFP fusion retains antigenic and receptor-binding activity.

### 4.7 Future Directions

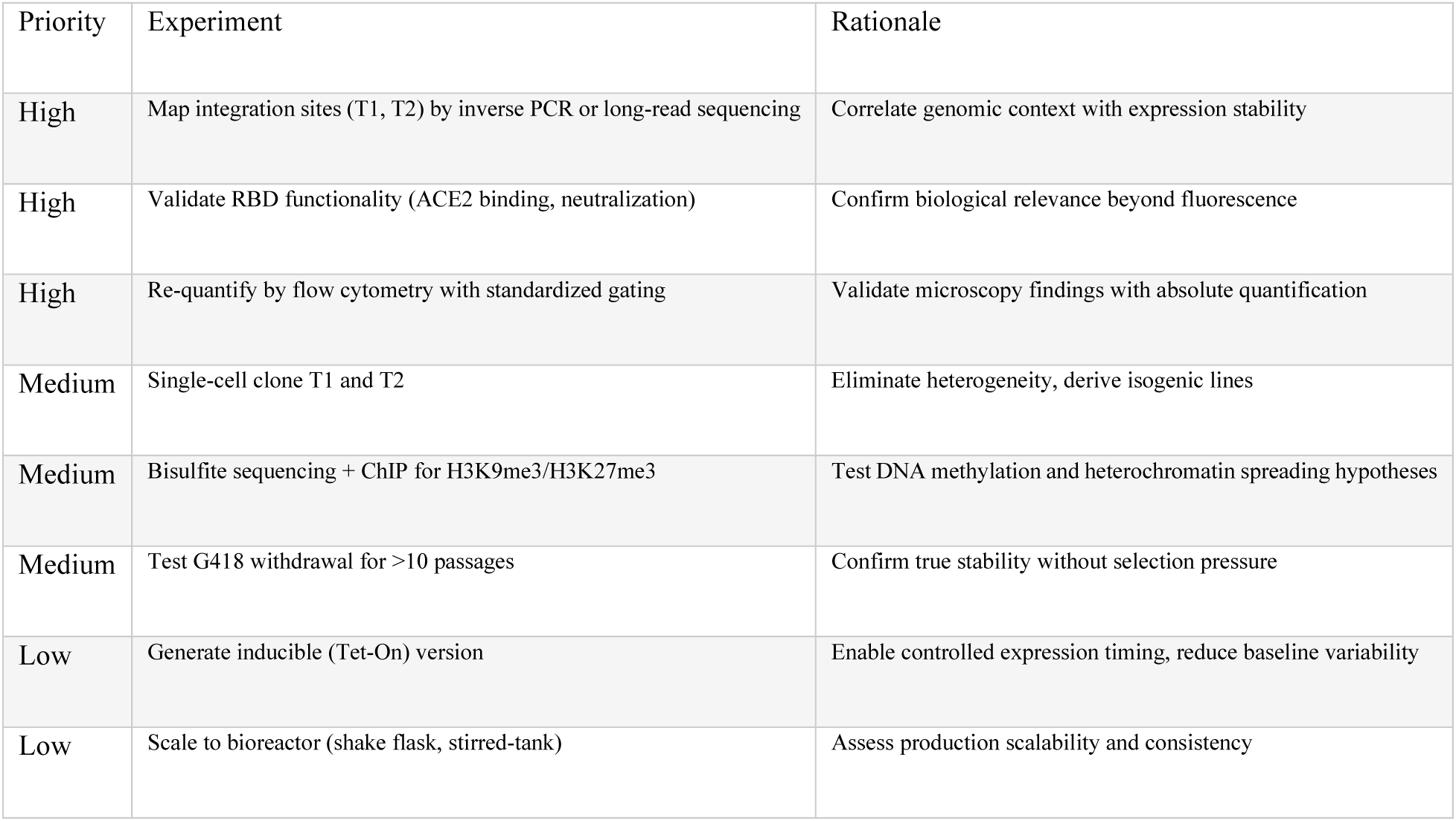

| Priority | Experiment | Rationale |
| --- | --- | --- |
| High | Map integration sites (T1, T2) by inverse PCR or long-read sequencing | Correlate genomic context with expression stability |
| High | Validate RBD functionality (ACE2 binding, neutralization) | Confirm biological relevance beyond fluorescence |
| High | Re-quantify by flow cytometry with standardized gating | Validate microscopy findings with absolute quantification |
| Medium | Single-cell clone T1 and T2 | Eliminate heterogeneity, derive isogenic lines |
| Medium | Bisulfite sequencing + ChIP for H3K9me3/H3K27me3 | Test DNA methylation and heterochromatin spreading hypotheses |
| Medium | Test G418 withdrawal for >10 passages | Confirm true stability without selection pressure |
| Low | Generate inducible (Tet-On) version | Enable controlled expression timing, reduce baseline variability |
| Low | Scale to bioreactor (shake flask, stirred-tank) | Assess production scalability and consistency |

### 4.8 Conclusion

We have established and characterized HPSF-IM-RBD-BHSKPU T1-T4, G418-selected, immortalized human primary splenic fibroblast lines stably expressing SARS-CoV-2 RBD-sfGFP. The cells demonstrate stable transgene retention through cryopreservation and 98 days of continuous culture, confirming chromosomal integration. However, expression is not static -- it exhibits passage-dependent oscillation and progressively increasing heterogeneity consistent with position-effect variegation. The divergence among T1-T4 highlights the critical role of integration site context in determining transgene behavior. These findings establish HPSF-IM-RBD-BHSKPU as both a practical platform for RBD production (optimal at P1-P4) and a model system for studying epigenetic regulation of transgene expression in human primary cell backgrounds. Future molecular characterization of integration sites and epigenetic marks will transform these descriptive observations into mechanistic understanding, with direct implications for rational cell line engineering in biopharmaceutical manufacturing.

## Acknowledgments

This work is the result of wonderful community support at Kwantlen Polytechnic University (KPU). We owe our sincere thanks to Dr. Brett Favaro (Dean of Science), Christina Heinrick (Associate Dean of Science), and Jackie Au (Research Services) for making the administrative process of naming our new cell lines (HPSF-IM-RBD-BHSKPU1 and HPSF-IM-RBD- BHSKPU2) smooth and straightforward. We are grateful to Christina Iggulden and David Robinson for their thorough lab safety training and assistance in the laboratory. Our deepest gratitude goes to Dr. Barnabe D. Assogba for supervising this project; his steady guidance, constant encouragement, and trust in our potential as emerging scientists kept us motivated through every challenge. Finally, we want to thank the Office of Research Services. The Student Research and Innovation Grant (SRIG) provided vital financial support, allowing us to fully immerse ourselves in our research and successfully achieve our goals.

## Supplementary Materials

Not applicable

## Author Contributions

K. S. Maan: Conceptualization, Methodology, Investigation, Data Collection, Funding acquisition, Writing – original draft. Z. A. Baloch: Data Curation, Visualization. I. Vashishat: Data Curation, Visualization. S. S. Bhullar: Data Curation, Visualization. B. D. Assogba: Conceptualization, Methodology, Resources, Project administration, Supervision, Funding acquisition, Writing, review & editing.

## Funding

This work was funded by Kwantlen Polytechnic University Student Research & Innovation Grant, under grant #

## Institutional Review Board Statement

Not applicable.

## Informed Consent Statement

Not applicable

## Conflicts of Interest

The authors declare no conflicts of interest.

## Data Availability

All data generated during this study are included in this published article. Raw fluorescence data, normalized values, and analysis scripts are available within the supplementary materials. The primary data file consists of an Excel workbook containing separate sheets for: (1) raw data, (2) normalized data, (3) summary statistics, (4) regression analysis, (5) coefficient of variation (CV) calculations, and (6) figure generation parameters. Raw microscopy images and ImageJ analysis scripts are available at Kwantlen Polytechnic University upon reasonable request.

## Notes

### Competing Interest Statement

The authors have declared no competing interest.

